# CoCl_2_ triggered pseudohypoxic stress induces proteasomal degradation of SIRT4 *via* polyubiquitination of lysines K78 and K299

**DOI:** 10.1101/2023.05.16.540489

**Authors:** Nils Hampel, Jacqueline Georgy, Mehrnaz Mehrabipour, Alexander Lang, Jürgen Scheller, Mohammad R. Ahmadian, Doreen M. Floss, Roland P. Piekorz

**Author notes:** Present address: Department of Cardiology, Pulmonology, and Vascular Medicine, Medical Faculty, Heinrich Heine University Düsseldorf, Germany (Alexander Lang).

## Abstract

SIRT4 comprises together with SIRT3 and SIRT5 the mitochondrially localized subgroup of sirtuins. SIRT4 regulates *via* its NAD^+^-dependent enzymatic activities mitochondrial bioenergetics, dynamics (mitochondrial fusion), and quality control (mitophagy). Here, we address the regulation of SIRT4 itself by characterizing its protein stability and degradation upon CoCl_2_-induced pseudohypoxic stress that typically triggers mitophagy. Interestingly, within the mitochondrial sirtuins, only the protein levels of SIRT4 or ectopically expressed SIRT4-eGFP decrease upon CoCl_2_ treatment of HEK293 cells. Co-treatment with BafA1, an inhibitor of autophagosome-lysosome fusion required for autophagy/mitophagy, or the use of the proteasome inhibitor MG132 prevented CoCl2-induced SIRT4 downregulation. Consistent with the proteasomal degradation of SIRT4, the lysine mutants SIRT4(K78R) and SIRT4(K299R) showed significantly reduced polyubiquitination upon CoCl_2_ treatment and were more resistant to pseudohypoxia-induced degradation as compared to SIRT4. Moreover, SIRT4(K78R) and SIRT4(K299R) displayed increased basal protein stability as compared to wild-type SIRT4 when subjected to MG132 treatment or cycloheximide (CHX) chase assays. Thus, our data indicate that stress-induced protein degradation of SIRT4 occurs through two mechanisms, (i) *via* mitochondrial autophagy/mitophagy, and (ii) as a separate process *via* proteasomal degradation within the cytoplasm.

## 1. Introduction

Sirtuins comprise a group of proteins initially defined through the identification of the NAD^+^- dependent histone deacetylase Sir2 in yeast [1]. Sirtuins can be sub-divided into five distinct phylogenetic branches by analysis of conserved catalytic core domain sequences. In human, seven sirtuins have been identified grouping them into four phylogentic branches, *i.e.,* class 1 (sirtuins 1 to 3), class 2 (SIRT4), class 3 (SIRT5), and class 4 (sirtuins 6 and 7) [2,3]. These proteins function in epigenetic regulation and gene expression control in the nucleus (SIRT1, 2, 6, and 7; [4]), microtubule dynamics (SIRT2, SIRT4; [5-7]), proliferation/cell survival, senescence and ageing (*e.g.* SIRT4 and SIRT6; [8,9]), life-span regulation (*e.g.* SIRT6; [9,10]), and regulation of mitochondrial metabolism (SIRT3, 4, 5; [11,12]). Mitochondrial sirtuins like SIRT3 represent potential targets for the treatment of aging-associated diseases [13,14]. This is further emphasized by recent data indicating an involvement of SIRT4 in the onset and development of Parkinson’s disease [15].

Human sirtuins localize in multiple subcellular compartments, functioning across them [16-20]. *E.g*., SIRT4 is distributed between the cytoplasm, nucleus, and in particular mitochondria [5,21], the latter based on an N-terminal mitochondrial targeting sequence typical for mitochondrial sirtuins [22-24]. Functionally, SIRT4 has been characterized in mitochondria as tumor suppressor and inhibitor of the metabolic gatekeeper enzymes pyruvate dehydrogenase (PDH; [25]) and glutamate dehydrogenase (GDH; [26,27]) as well as based on its deacetylase activity as a regulator of leucin metabolism and insulin secretion [28]. Moreover, several recent reports attributed novel extramitochondrial roles to SIRT4 in microtubule dynamics and regulation of mitotic cell cycle progression, WNT/β-Catenin and Hippo signaling, and SNARE complex formation required for autophagosome-lysosome fusion [5,29-31].

The expression of SIRT4 is regulated both at the gene/mRNA and protein level. Regarding the latter, the degradation of sirtuins is mediated by two major cellular pathways, macroautophagy and presumably the ubiquitin-proteasome pathway. Mitochondrially localized sirtuins are degraded by macroautophagy in neuronal LUHMES cells, a M. Parkinson disease model, upon MPP^+^ (1-methyl-4-phenylpyridinium) induced oxidative stress [32]. This degradation of oxidized sirtuins could be prevented by treatment with Bafilomycin A1 (BafA1), an inhibitor of autophagosome-lysosome fusion and therefore (macro)autophagy, whereas treatment with MG132, a widely used proteasome inhibitor, failed to preclude reduction of sirtuin protein levels [32].

Interestingly, within human SIRT4 comprehensive proteome mapping identified the putative Ubiquitin target lysine residues K78 and K299 [33,34], thus indicating that SIRT4 may indeed undergo ubiquitination and possibly polyubiquitination, given its subcellular distribution not only in mitochondria, but also in the cytoplasm and nucleus [5,21]. Polyubiquitination occurs *via* the internal lysine residue K48 of Ubiquitin (K48-polyUb), that is required to tag target proteins by multiple Ubiquitin molecules for subsequent proteasomal degradation in the cytoplasm [35-37]. Interestingly, hypoxia leads to downregulation of SIRT4 at the protein level [38,39] by unknown mechanism(s). Therefore, in the present study we employed a chemical hypoxia model using CoCl_2_ treatment [40] to address the role of the SIRT4 lysine residues K78 and K299 [33,34] in basal protein stability and stress-induced polyubiquitination and proteasomal degradation of SIRT4.

## 2. Materials and Methods

### 2.1 Reagents

CoCl_2_, Bafilomycin A1, Cycloheximide (CHX), and MG132 were obtained from Roth, Cayman Chemical, Sigma-Aldrich, and Selleck Chemicals, respectively. Primary antibodies were directed against SIRT3 (Cell Signaling #5490), SIRT4 (Proteintech #66543-1-Ig), SIRT5 (Cell Signaling #8782), eGFP (Roche #11814460001), Ubiquitin (Proteintech #12986-1-AP; Cell Signaling #3933), and α- Tubulin (Abcam #ab52866, Proteintech #11224-1-AP). Primary antibodies were detected using anti-mouse (700 nm; LI-COR IRDye #926-32213) or anti-rabbit (800 nm; LI-COR IRDye #926-68072) secondary antibodies.

### 2.2 Cell culture

Parental and SIRT4 wild-type/mutant expressing HEK293 cell lines were cultured at 37°C and 5% CO_2_ in DMEM (Dulbecco’s Modified Eagle Medium) containing high glucose (4.5 g/L; Thermo Fisher Scientific) with 10% fetal bovine serum (FBS; Gibco) and penicillin (100 units/mL)/streptomycin (100 μg/mL) (Genaxxon). HEK293 cells were obtained from the German Collection of Microorganisms and Cell Cultures GmbH (DSMZ) (HEK293: ACC 305). HEK293-eGFP and HEK293-SIRT4-eGFP cell lines have been described previously [5,41].

### 2.3 Site-directed mutagenesis

Primers to generate SIRT4 mutations K78R and K299R were obtained from Eurofins Genomics. The sequences of the oligonucleotides used in this study will be provided upon request. The pcDNA3.1 vector containing SIRT4-eGFP was used as a template for PCR-based site-directed mutagenesis using 100 picomoles of forward and reverse primers, 10-20 ng of template plasmid, and 1 µL of Phusion High-Fidelity DNA Polymerase (Thermo Fisher Scientific). PCR reactions were performed for 15 cycles at a denaturation temperature of 98°C (1 min), an annealing temperature of 55°C (1 min), and an extension temperature of 72°C (3 min). Methylated template DNA was digested by DpnI afterwards. SIRT4 point mutations were confirmed by sequencing (MicroSynth Seqlab GmbH).

### 2.4 Generation of SIRT4 expressing cell lines

HEK293 cell lines stably expressing the mutated SIRT4-eGFP variants (K78R, K299R, or K78R/K299R) have been generated using the Turbofect transfection reagent (ThermoFisher Scientific) and cultured in media containing 400 µg/ml Geneticin/G418 (Genaxxon) as a permanent selection agent. The expression of SIRT4-eGFP fusion constructs was validated by immunoblotting and flow cytometry. Generation of HEK293-SIRT(H161Y)-eGFP and HEK293-SIRT4(ΔN28)-eGFP cell lines has been previously described [5,41].

### 2.5 Treatment of HEK293 cell lines with the pseudohypoxia agent CoCl_2_

HEK293 cell lines were grown to a cell density of approximately 80% and then subjected to a chemical hypoxia model [40] using CoCl_2_ treatment at concentrations of 250 µM and 400 µM for 24 h or 36 h.

### 2.6 Pulse-chase protein stability assay using Cycloheximide

To determine basal protein stability of SIRT4 and mutants thereof, HEK293 cell lines were treated at a cell density of approximately 80% with the protein biosynthesis inhibitor Cycloheximide for 4 h, 8 h, and 24 h. Based on this chase kinetics, linear regression was employed to calculate the protein half-life of SIRT4 variants.

### 2.7 Preparation of total cell lysates and immunoblot analysis

Cleared cell lysates were generated using lysis buffer containing 0.3% CHAPS (3-[(3- Cholamidopropyl) dimethylammonio]-1-propanesulfonate), 50 mM Tris-HCl (pH 7.4), 150 mM NaCl, 1 mM Na_3_VO_4_, 10 mM NaF, 1 mM EDTA, 1 mM EGTA, 2.5 mM Na_4_O_7_P_2_, and 1 µM DTT. The cOmplete™ protease inhibitor cocktail (Sigma-Aldrich) was used to prevent the degradation of proteins in the lysates. The latter were cleared by centrifugation (11.000 x g at 4°C for 20 min) and the protein concentration of the supernatants (total cell lysates) was determined using the Bradford assay (Roth). Relative quantification of protein levels (as compared to α-Tubulin or β-Actin loading controls) was performed by ImageJ-based densitometric analysis of specific immunoblot signals.

### 2.8 Immunoprecipitation of ubiquitinated SIRT4-eGFP wild-type and mutant proteins

Total cell lysates were obtained as described above and subjected to immunoprecipitation analysis using single-domain anti-eGFP antibodies (nanobody method based on [42]) essentially as described [5,41]. Polyubiquitination of wild-type and mutant SIRT4-eGFP forms was detected using Ubiquitin-specific antibodies.

### 2.9 Phylogenetic analysis

Sequences were obtained from the UniProt database (www.uniprot.org) and further analysis was performed using the ClustalW multiple alignment method followed by the sequence alignment editor software BioEdit 7.2.5.

### 2.10 Statistical analysis

Data are presented as mean ± S.D. Multiple comparisons were analyzed by one-way or two-way analysis of variance (ANOVA) followed by Tukey’s post-hoc test to identify group differences in variance analysis using the GraphPad Prism software. Statistical significance was set at the level of p ≤ 0.05 (*p ≤ 0.05, **p ≤ 0.01, ***p ≤ 0.001).

## 3. Results

### 3.1 The protein levels of SIRT4, but not SIRT3 and SIRT5, decrease upon induction of pseudohypoxia

Given that hypoxia leads to downregulation of SIRT4 at the protein level [38,39], we tested the impact of CoCl_2_-induced pseudohypoxia on all three mitochondrial sirtuins in HEK293 cells. As shown in Figure 1, CoCl_2_ treatment at concentrations of 250 µM and 400 µM for 24 h resulted in a decrease of endogenous SIRT4 protein levels by up to 50%. In contrast, under the same conditions, total cell protein quantities of SIRT3 and SIRT5 did not alter significantly. Thus, within the family of mitochondrial sirtuins, the expression of SIRT4 is specifically downregulated at the protein level upon pseudohypoxic stress, presumably independent of altered SIRT4 gene expression as evident from the study by Pecher et al. [39].

**Figure 1.**
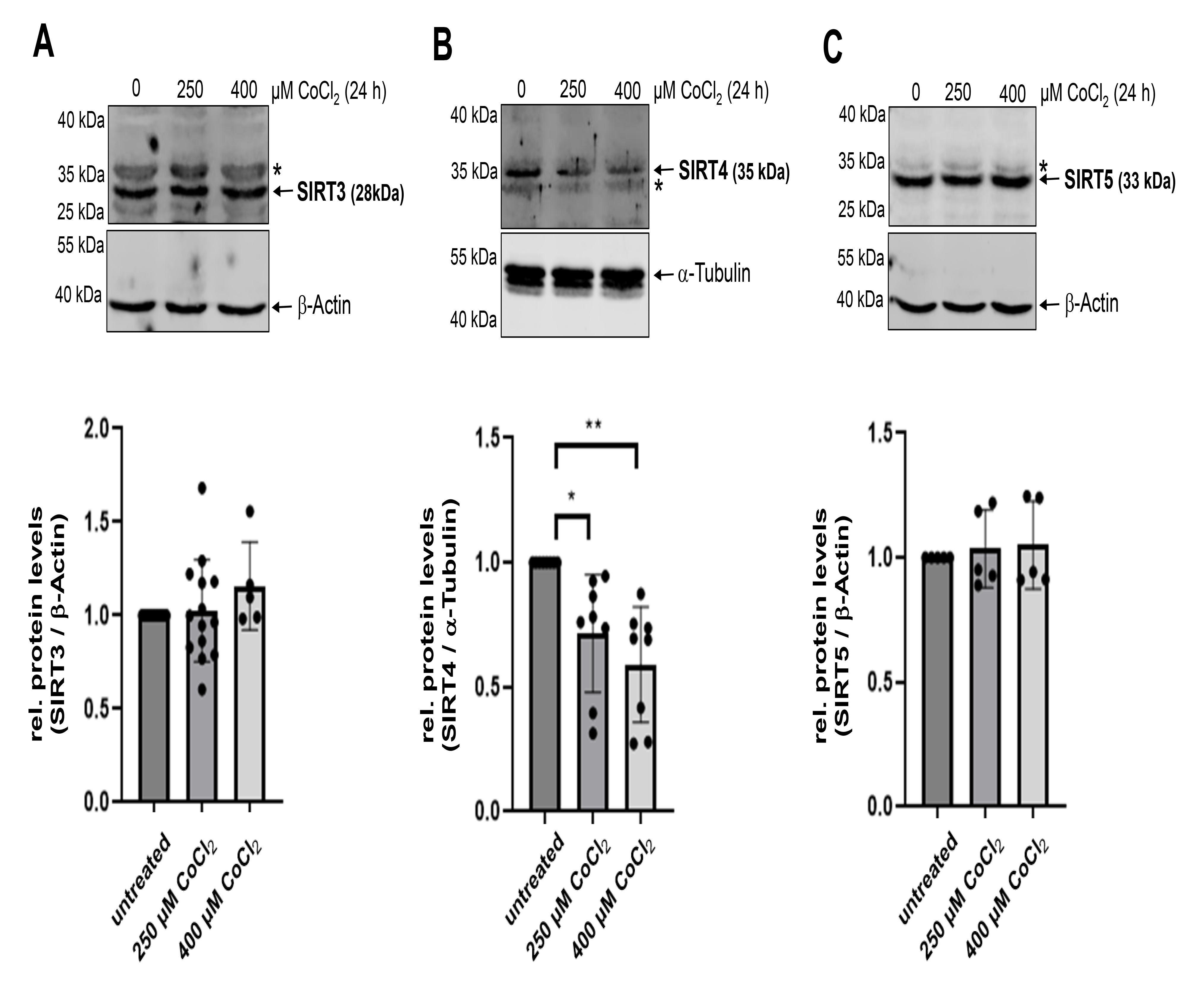
Protein levels of SIRT4, but not SIRT3 and SIRT5, are downregulated upon CoCl2-induced pseudohypoxia. HEK293 cells were subjected to CoCl_2_ treatment for 24 h followed by analysis of endogenous protein levels of SIRT3 (**A**), SIRT4 (**B**), and SIRT5 (**C**). Relative quantification of immunoblot signals was performed using ImageJ-based densitometric evaluation and α-Tubulin levels as loading control. Unspecific bands are marked (*). To determine statistical significance, a One-Way ANOVA test followed by Tukeýs test was employed (mean ± S.D.; * *p* < 0.05; ** *p* < 0.01).

### 3.2 Inhibition of the proteasome or autophagic degradation prevents protein degradation of SIRT4 in CoCl_2_ induced pseudohypoxia

Consistent with the findings for endogenous SIRT4 (Figure 1), the protein levels of ectopically expressed SIRT4-eGFP (Figure 2A,B), but not eGFP as control (Figure S1), were also reduced by approximately 60% upon CoCl_2_-treatment. Interestingly, this reduction of SIRT4-eGFP levels could be prevented by co-treatment with either BafA1 or MG132, indicating that both macroautophagy/mitophagy and the proteasome, respectively, are involved in pseudohypoxic stress induced SIRT4 degradation. MG132 mediated inhibition of the proteasome led also to the stabilization of the catalytically inactive mutant SIRT4(H161Y) and the N-terminal deletion mutant SIRT4(ΔN28) that lacks the mitochondrial translocation sequence (Figure S2). Thus, proteasomal degradation of SIRT4 is independent of its enzymatic activity and occurs extra-mitochondrially in the cytoplasm.

**Figure 2:**
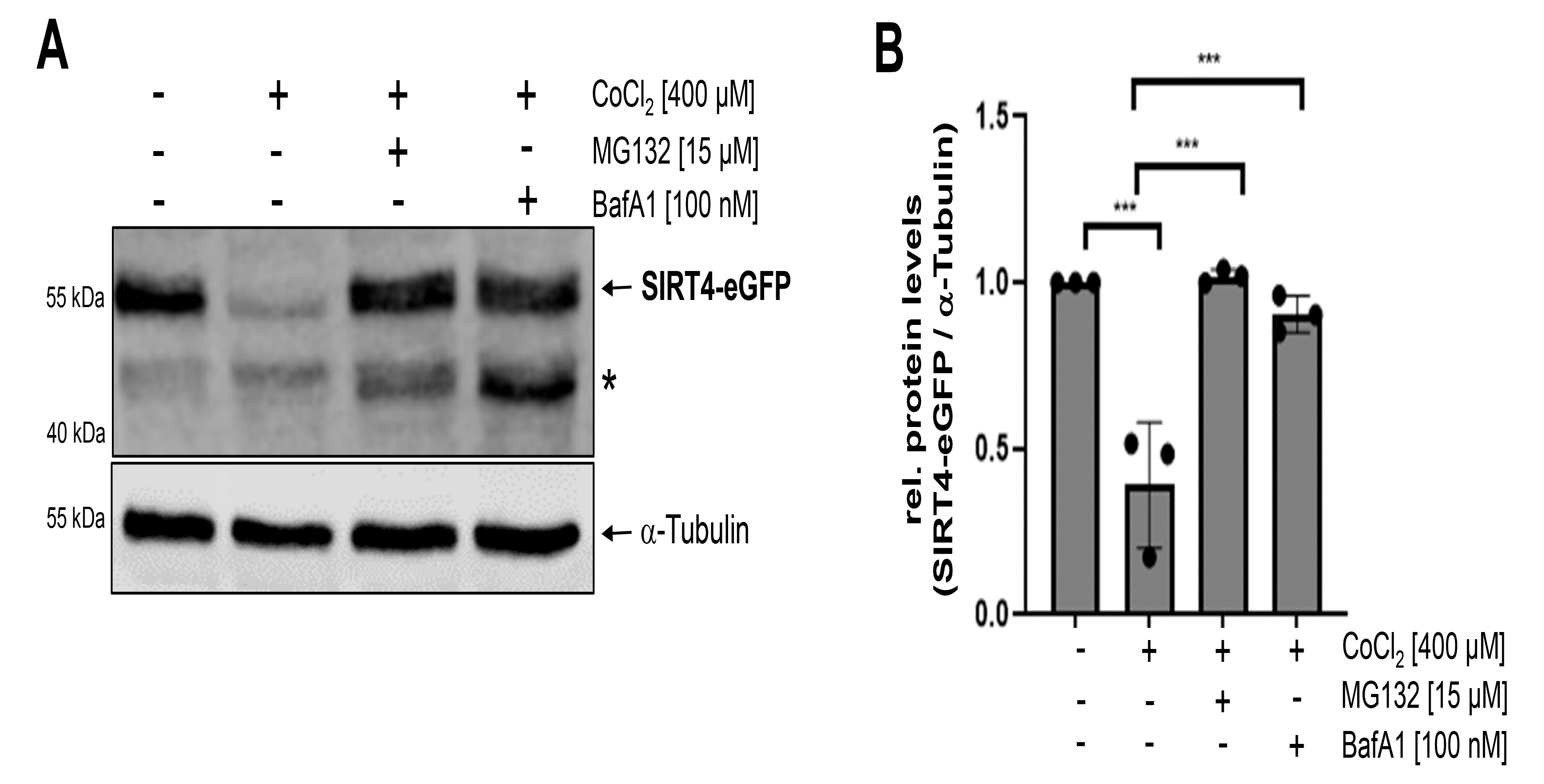
Co-treatment of HEK293-SIRT4-eGFP cells with MG132 or Bafilomycin A1 (BafA1) inhibits degradation of SIRT4-eGFP during CoCl_2_-induced pseudohypoxia. SIRT4-eGFP expressing HEK293 cells were subjected to CoCl_2_ treatment for 24 h in the presence or absence of the proteasome inhibitor MG132 or BafA1, which prevents autophagosome-lysosome fusion. SIRT4-eGFP protein levels were analyzed by immunoblotting using anti-eGFP antibodies (**A**) and ImageJ-based densitometric evaluation using α-Tubulin levels as loading control (**B**). Unspecific bands are marked (*). To determine statistical significance, a One-Way ANOVA test followed by Tukeýs test was employed (mean ± S.D.; *** *p* < 0.001).

### 3.3 The SIRT4 lysine mutants K78R and K299R are stabilized in CoCl_2_ induced pseudohypoxia

Proteome-wide mapping identified within human SIRT4 the putative Ubiquitin target lysine residues K78 and K299 [33,34]. Thus, to further characterize the role of ubiquitination and proteasomal degradation in stress-induced regulation of SIRT4 levels, we generated HEK293 cell lines stably expressing the Lysine to Arginine mutated SIRT4 variants K78R, K299R, or the double mutant K78R/K299R (Figure S3), therefore preventing ubiquitination of these lysine residues. We subjected these cell lines to CoCl_2_ induced pseudohypoxic stress followed by analysis of wild-type and mutated SIRT4 protein levels. As indicated in Figure 3, CoCl_2_ treatment for 24 h resulted in a significant reduction of SIRT4-eGFP protein levels by approximately 50%, whereas the mutants K78R, K299R, and K78R/K299R were stable with no overtly quantitative changes. Longer CoCl_2_ treatment for 36 h ameliorated this phenotype and resulted in significant degradation of all three mutants, although K299R still retained an increased stability. Thus, both lysine residues K78 and K299 regulate the protein stability of SIRT4 upon pseudohypoxic stress.

**Figure 3.**
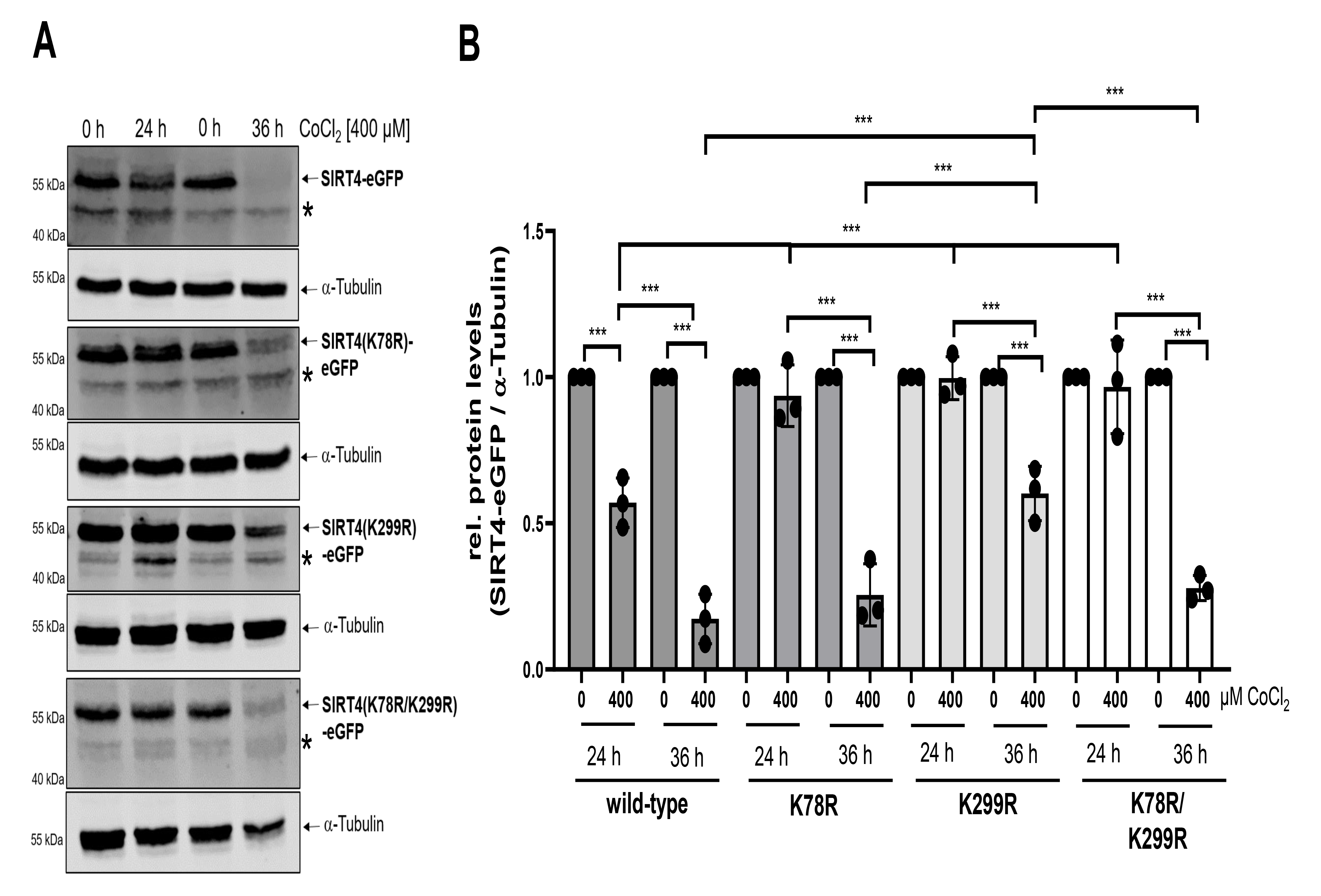
The SIRT4 mutants K78R and K299R are more resistant to CoCl_2_-induced degradation. HEK293 cell lines expressing SIRT4-eGFP or the indicated SIRT4 mutants thereof were subjected to CoCl_2_-induced pseudohypoxia for 24 h and 36 h followed by immunoblot analysis of the respective SIRT4-eGFP/mutant SIRT4-eGFP levels using anti-eGFP antibodies (**A**) and ImageJ based densitometric evaluation using α-Tubulin levels as loading control (**B**). Unspecific bands are marked (*). To determine statistical significance, a Two-Way ANOVA test followed by Tukeýs test was employed (mean ± S.D.; *** *p* < 0.001).

### 3.4 The SIRT4 lysine mutants K78R and K299R undergo decreased polyubiquitination upon CoCl_2_ induced pseudohypoxia

Polyubiquitination (poly-Ub) functions as a precursor and initiator of proteasome-mediated protein degradation [43]. We next subjected wild-type and mutated SIRT4-eGFP from untreated and CoCl_2_- treated cells to immunoprecipitation using anti-eGFP nanobody beads followed by the analysis of the degree of SIRT4 polyubiquitination using anti-Ubiquitin immunoblotting. Consistent with the previous findings, the stress-induced polyubiquitination of all three SIRT4 variants K78R, K299R, and K78R/K299R was significantly lower as compared to wild-type SIRT4, the latter showing a 3-fold induction in poly-Ub levels (Figure 4A,B). Next, we explored the conservation of lysine residues K78 and K299 of human SIRT4 both within the mammalian sirtuins and evolutionary within known SIRT4 homologues. As indicated in Figure 4C, both K78 and K299 are unique for SIRT4 among all seven human sirtuin family members, in particular the mitochondrial sirtuins. The only exception is K299 which is also found in all known SIRT1 isoforms, but this lysine residue does not seem to be involved in SIRT1 ubiquitination [44]. At the level of SIRT4 homologs, K78 seems highly conserved in mammals but is absent in phylogenetically more distant species like *Xenopus tropicalis* (Figure 4D). In contrast, lysine K299 is completely conserved throughout the vertebrates indicating that K299 plays an evolutionary more conserved role in the regulation of stress-induced proteasomal degradation of SIRT4. Overall, these findings identify the SIRT4 residues K78 and K299 as conserved polyubiquitination targets and indicate that the level of polyubiquitination of SIRT4 negatively correlates with its protein stability upon pseudohypoxic stress.

**Figure 4.**
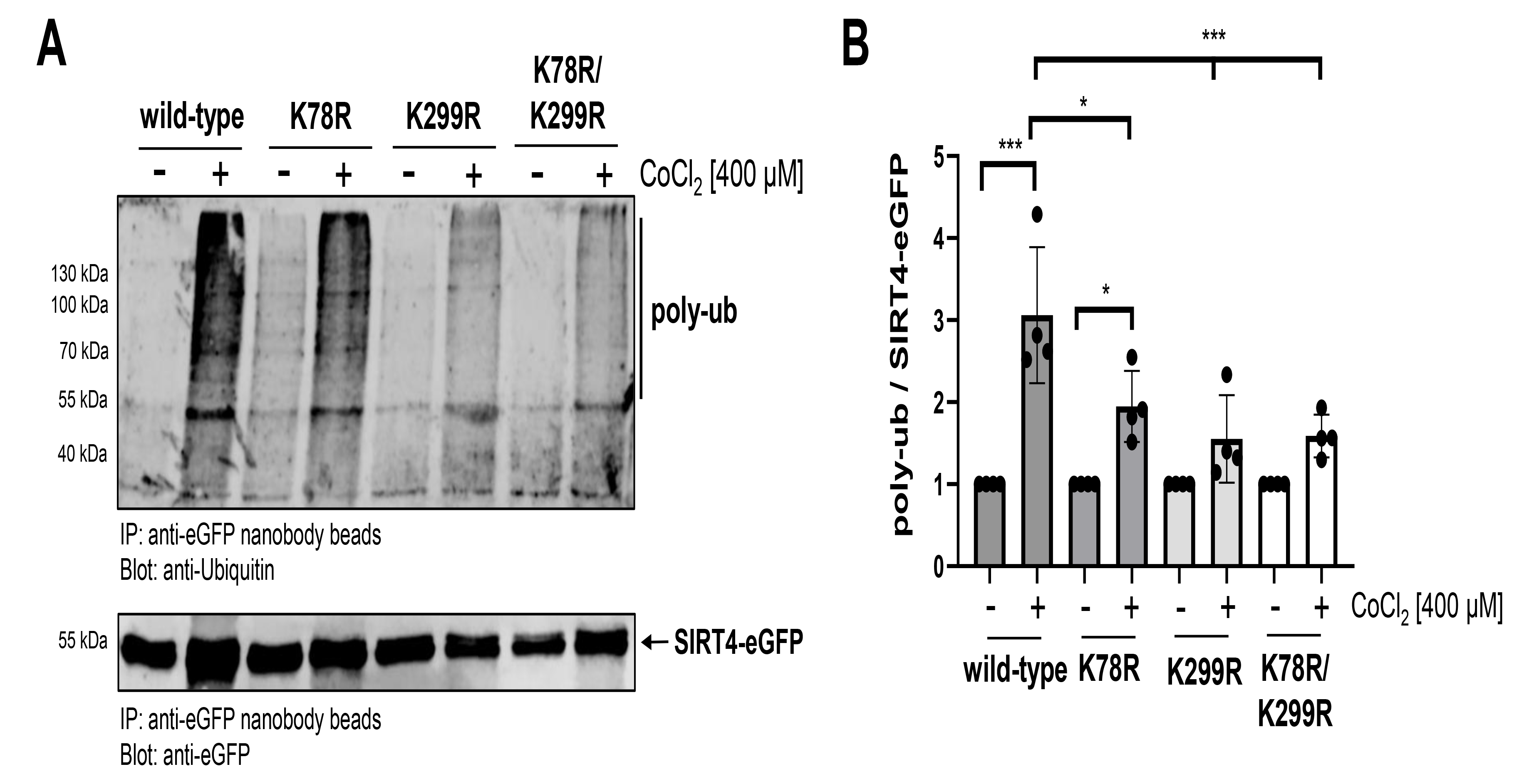
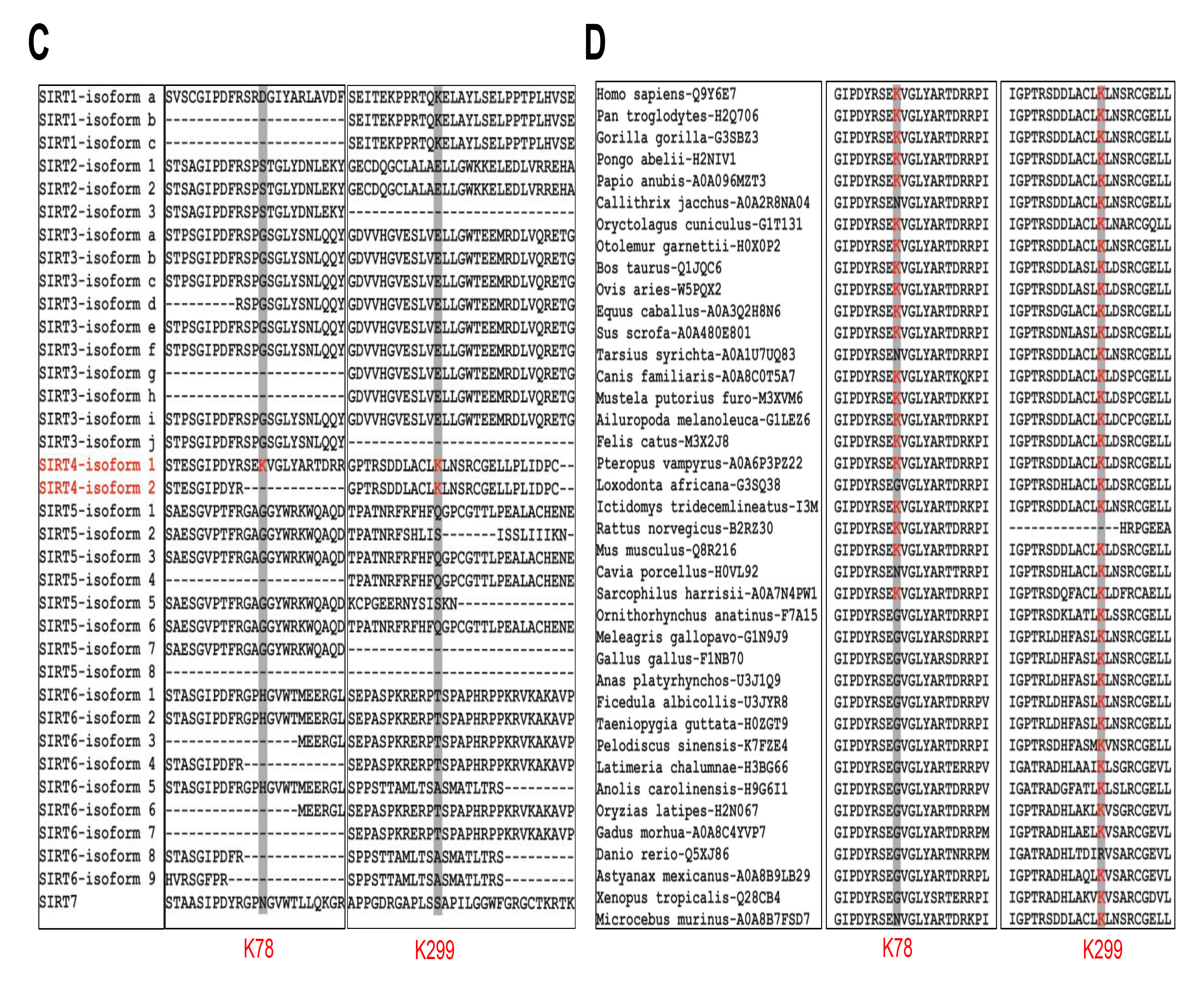
The SIRT4 mutants K78R and K299R show a decreased polyubiquitination upon CoCl_2_- induced pseudohypoxia. (**A**) HEK293 cell lines expressing SIRT4-eGFP or the indicated SIRT4-eGFP mutants thereof were either untreated or subjected to CoCl_2_-induced pseudohypoxia for 24 h. SIRT4 variants were immunoprecipitated using anti-eGFP nanobody beads and further analyzed for the degree of polyubiquitination (poly-ub) using anti-Ubiquitin immunoblotting (upper panel). Immunoprecipitated SIRT4-eGFP proteins were detected on the same membrane using anti-eGFP antibodies (lower panel). (**B**) Relative quantification of polyubiquitination of SIRT4-eGFP mutants compared to wild-type SIRT4-eGFP using ImageJ-based densitometric evaluation. To determine statistical significance, a Two-Way ANOVA test followed by Tukeýs test was employed (mean ± S.D.; * *p* < 0.05; *** *p* < 0.001). (**C**) SIRT4-specific conservation of lysines K78 and K299 (marked in red) within the human Sirtuin protein family. (**D**) Analysis of evolutionary conservation of K78 and K299 (marked in red) in SIRT4 homologues of vertebrates. Sequences in (**C**) and (**D**) were obtained from the UniProt database (www.uniprot.org). Sequence analysis was performed using the ClustalW multiple aligment method followed by the sequence alignment editor software BioEdit 7.2.5.

### 3.5 The SIRT4 lysine mutants K78R and K299R display an increased basal protein stability

Pulse chase assays are established to analyze the degree of basal protein stability upon cycloheximide (CHX) mediated inhibition of protein translation [45]. Therefore, HEK293 cell lines expressing wild-type SIRT4 or the mutants K78R, K299R, or K78R/K299R, were subjected to a time kinetics of CHX treatment for up to 24 h. Interestingly, all three mutants displayed a delayed decrease in protein levels as compared to wild-type SIRT4 (Figure 5A,B). To further examine differences in stability between wild-type SIRT4 and its mutants we calculated their protein half-lives (T_1/2_) (Figure 5C). The T_1/2_ for SIRT4(K78R) was approximately 1.6-fold increased as compared to wild-type SIRT4, whereas its difference to SIRT4(K299R) was nearly significant. To address whether these SIRT4 mutants are more resistant to proteasomal degradation under basal conditions, we treated SIRT4 wild-type/mutant-expressing cell lines for 24 h with MG132. As indicated in Figures 5A and 5D, in contrast to wild-type SIRT4, all SIRT4 variants showed a clear increase in protein levels upon MG132 treatment, with the biggest significant effect on the double mutant K78/K299.

**Figure 5.**
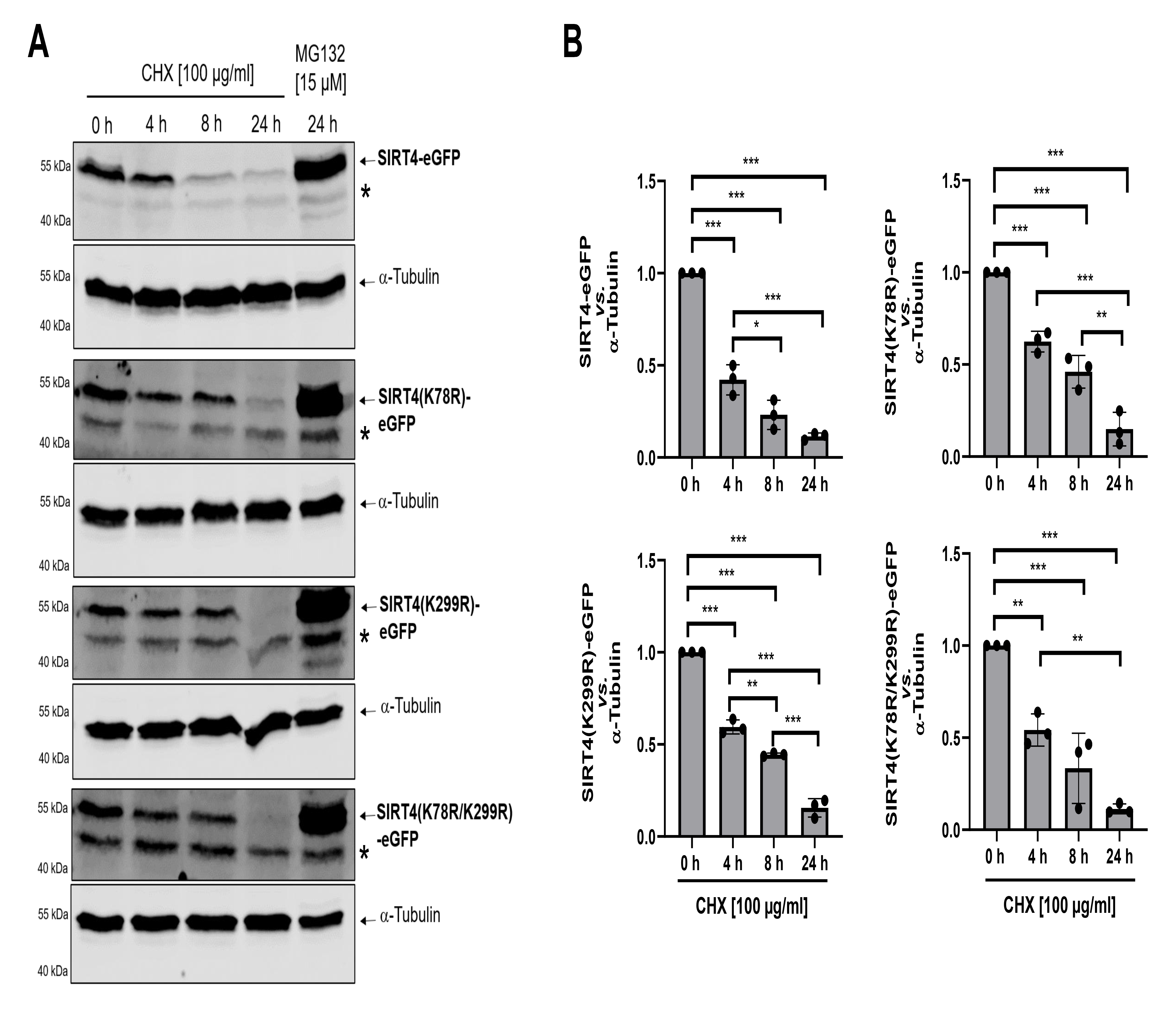
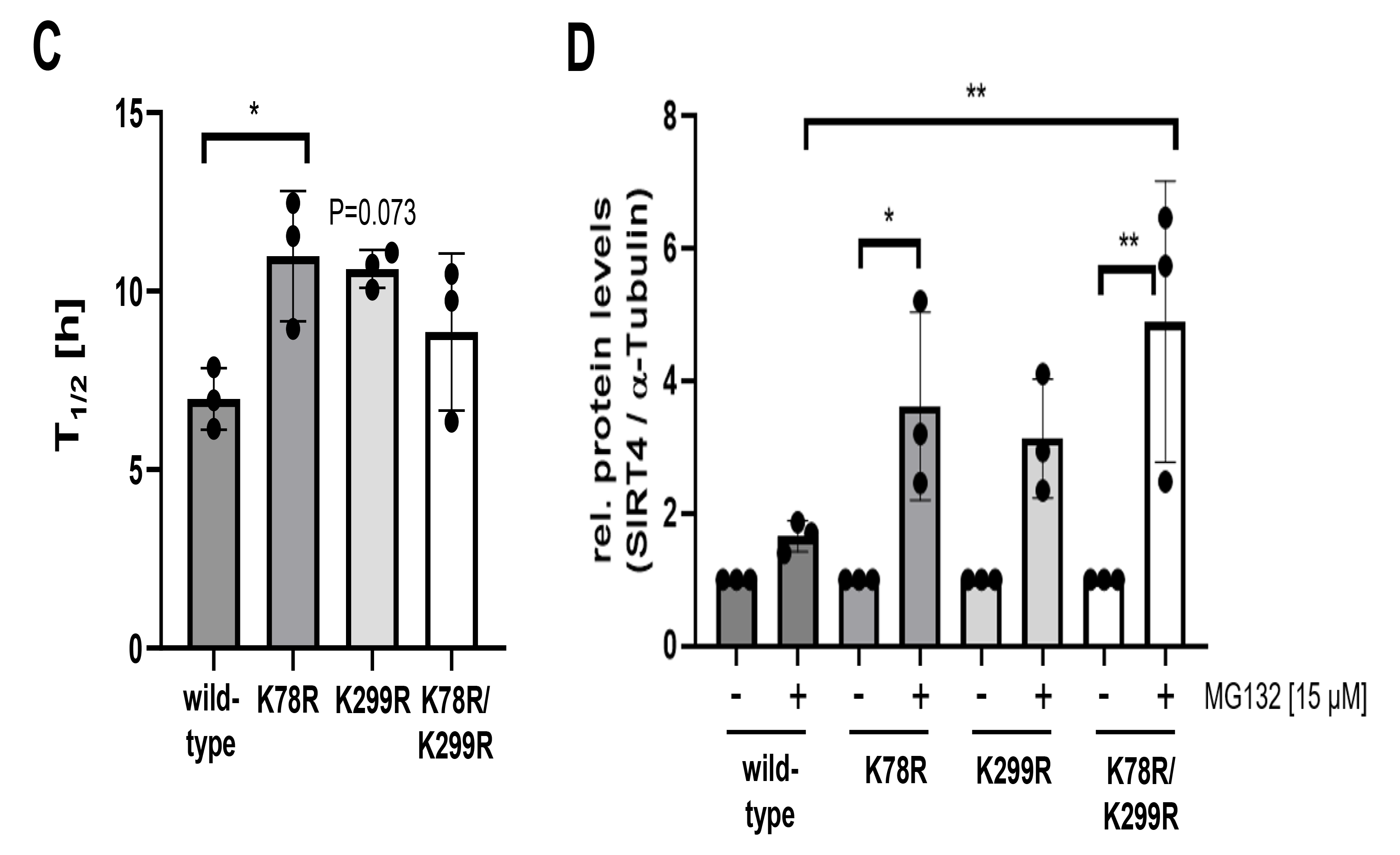
The SIRT4 mutants K78R and K299R display increased protein stability. HEK293 cells stably expressing SIRT4-eGFP or the indicated SIRT4-eGFP mutants thereof were subjected to cycloheximide (CHX) chase assays for 4 h, 8 h, and 24 h or treatment with the proteasome inhibitor MG132 for 24 h. Expression of SIRT4-eGFP or the SIRT4-eGFP mutants thereof was analyzed by immunoblotting using anti-eGFP antibodies (**A**) and ImageJ-based densitometric evaluation using α- Tubulin levels as loading control (**B**). Unspecific bands are marked (*). (**C**) Determination of the protein half-life (T_1/2_) of SIRT4-eGFP as compared to SIRT4(K78R)-eGFP and SIRT4(K299R)-eGFP analysed in CHX chase assays. (**D**) MG132 mediated inhibition of the proteasome increases the stability of SIRT4(K78R)-eGFP, SIRT4(K299R)-eGFP, and the double mutant SIRT4(K78R/K299R)- eGFP as compared to wild-type SIRT4-eGFP. Immunoblots were subjected to ImageJ-based densitometric evaluation using α-Tubulin levels as loading control. To determine statistical significance, Two-Way ANOVA tests followed by Tukeýs tests were employed (mean ± S.D.; * *p* < 0.05; ** *p* < 0.01; *** *p* < 0.001).

## 4. Discussion

This study provides novel insight into the posttranscriptional regulation of SIRT4 protein levels under stress, *i.e.,* pseudohypoxia-induced conditions. Our findings indicate that (i) pseudohypoxia-induced degradation of SIRT4 is mediated by two mechanisms, *via* macroautophagy/mitophagy upon mitochondrial translocation of SIRT4, and moreover as a separate process *via* proteasomal degradation within the cytoplasm; (ii) the latter mechanism depends on two conserved polyubiquitination targets of SIRT4, *i.e*., lysine residues K78 and K299; and (iii) within the group of mitochondrial sirtuins, SIRT4 is the only sirtuin which protein levels decrease upon CoCl_2_ induced pseudohypoxic stress. Consistent with this, downregulation of SIRT4 by hypoxia (1-2% O_2_) at the protein level occurs also under more physiological conditions in H9c2 cardio-myoblast and endothelial HUVEC cells [38,39]. This may result in an attenuated ROS response, given that increased SIRT4 can elevate mitochondrial H_2_O_2_ levels [41]. In contrast, dependent on the cell model analyzed, the modulation of SIRT3 by hypoxia results in either decrease or rather an increase of SIRT3 protein levels as summarized in [46]. *E.g*., a 2% O_2_ hypoxic condition leads to an increase of SIRT3 in endothelial HUVEC cells that preserves *via* deacetylation of FOXO3 bioenergetics and cell survival under hypoxia [47].

The expression of SIRT4 is regulated at both the gene/mRNA and protein level. At the transcriptional level, mTORC1 functions as a negative regulator by repressing *Sirt4* gene expression *via* degradation of the transcription factor CREB2 [48,49]. Moreover, the *Sirt4* gene is directly repressed by the lysine-specific demethylase 1 (Lsd1) [50]. Positive regulators of *Sirt4* gene expression include E2F1 [51], and interestingly also SIRT6, whose target genes *Sirt3* and *Sirt4* are downregulated upon SIRT6 deficiency resulting in mitochondrial dysfunction [52]. Lastly, several microRNAs (miR-15a-5p, miR-15b, miR- 130b-5p, and miR-497) bind SIRT4 transcripts and thereby modulate SIRT4 protein levels under basal as well as stress-induced and cell aging conditions [8,53-55].

The mechanism(s) involved in the direct protein degradation of mitochondrial sirtuins have only been recently addressed in closer detail by Baeken et al. [32]. The authors showed that MPP^+^ induced oxidative stress in neuronal LUHMES cells, a M. Parkinson disease model, results in the degradation of SIRT4. This could be prevented by treatment with BafA1, an inhibitor of autophagosome-lysosome fusion and therefore (macro)autophagy. Consistent with this, MPP^+^ treatment resulted in an increased sub-cellular co-localization of SIRT4 with LC3B positive autophagic structures [32]. In contrast, in the authors’ MPP^+^ model, the reduction of protein levels of oxidized SIRT4 was insensitive to treatment with the proteasome inhibitor MG132. This is in different from the rescue effect of MG132 treatment towards CoCl_2_-induced degradation of SIRT4 (Figure 2), and surprising, given that both MMP+, an inhibitor of mitochondrial complex I [56], and hypoxia, an inhibitor of complex III [57], lead to the accumulation of the mitochondrial ROS species H_2_O_2_. These contrary results could be based on the different cell models analyzed and/or due to different extents of ROS generated by MPP_+_ *vs.* CoCl_2_. In this regard, and given the dynamic subcellular distribution pattern of SIRT4 [5,21], one can speculate that lower to medium mitochondrial H_2_O_2_ levels target predominantly mitochondrially localized SIRT4, whereas high cellular H_2_O_2_ levels also lead to oxidation of cytosolically localized SIRT4. The latter would then require the proteasome besides (macro)autophagy for efficient SIRT4 degradation. However, it needs to be analysed to which extent CoCl_2_ treatment mediates SIRT4 degradation *via* ROS generation and subsequent SIRT4 oxidation.

Polyubiquitination of SIRT4 has been previously observed [58,59], but the ubiquitination site(s) of SIRT4 were not analysed. Consistent with our data, recent work by Zhao et al. identified lysine residue K78 of SIRT4 as a polyubiquitination target under basal, *i.e.,* non-stress conditions [60]. The authors’ data indicate that the mTORC1-c-Myc regulated E3-Ubiquitin protein ligase TRIM32 targets SIRT4 *via* polyubiquitination of lysine K78 for proteasomal degradation [60]. However, this mechanism may not be relevant under (pseudo)hypoxic conditions given that hypoxia downregulates TRIM32 protein levels as shown in pulmonary artery smooth muscle cells [61]. Thus, it remains to be determined (i) whether other SIRT4 interacting E3-Ubiquitin protein ligases, including RNF138 [60] or TRIM28 ([60] and own unpublished results), are involved in proteasomal degradation of SIRT4, and (ii) which of the lysine residues K78 and K299 are targeted by these E3- Ubiquitin ligases. Overall, our findings indicate that lysine K78 regulates protein half-life under basal conditions (Figure 5C), whereas polyubiquitination of lysine K299 mediates SIRT4 degradation upon cellular stress (Figures 3 and 4).

In eukaryotes, polyubiquitination-dependent proteasomal degradation of proteins takes place in the cytoplasm and in the nucleus [62]. Interestingly, recent findings identified a mitochondrial E3- ubiquitin ligase involved in the degradation of the mitophagy receptors BNIP3 and NIX [63] and further uncovered ubiquitin-dependent degradation of mitochondrial proteins at the inner mitochondrial membrane [64]. Given these observations one could speculate that polyubiquitination and proteasomal degradation of SIRT4 occurs not in the cytoplasm, but during/after mitochondrial translocation. Although we can exclude this possibility, the MG132-mediated stabilization of the ectopically expressed N-terminal deletion mutant SIRT4(ΔN28) (Figure S2), which cannot be imported into mitochondria, supports the existence of an extramitochondrial polyubiquitination and degradation mechanism for SIRT4.

## 5. Conclusions

We propose a model in which stress-induced degradation of SIRT4 is regulated by and dependent on its subcellular localization, *i.e*, macroautophagy of mitochondrially localized SIRT4 and the ubiquitin-proteasome mediated degradation of extra-mitochondrial/cytoplasmatic SIRT4 (Figure 6). Both degradation systems regulate cytoplasmatic *vs.* mitochondrial SIRT4 levels and therefore the respective subcellular functions of SIRT4. In the former case, SIRT4, a *bona fide* tumor suppressor protein, interacts with the mitotic spindle apparatus and negatively regulates cell cycle progression [5]. Here, downregulation of SIRT4 upon (pseudo)hypoxia would favor the proliferation of *e.g.* stem cells or tumor cells in hypoxic niches [65,66]. In the latter case, mitochondrial SIRT4 interacts with the GTPase OPA1 thereby favoring mitochondrial fusion and thus counteracting mitophagy [41,67]. Here, the downregulation of SIRT4 would promote mitophagy and prevent the accumulation of defective mitochondria due to hypoxia. These models need to be tested in the future.

**Figure 6.**
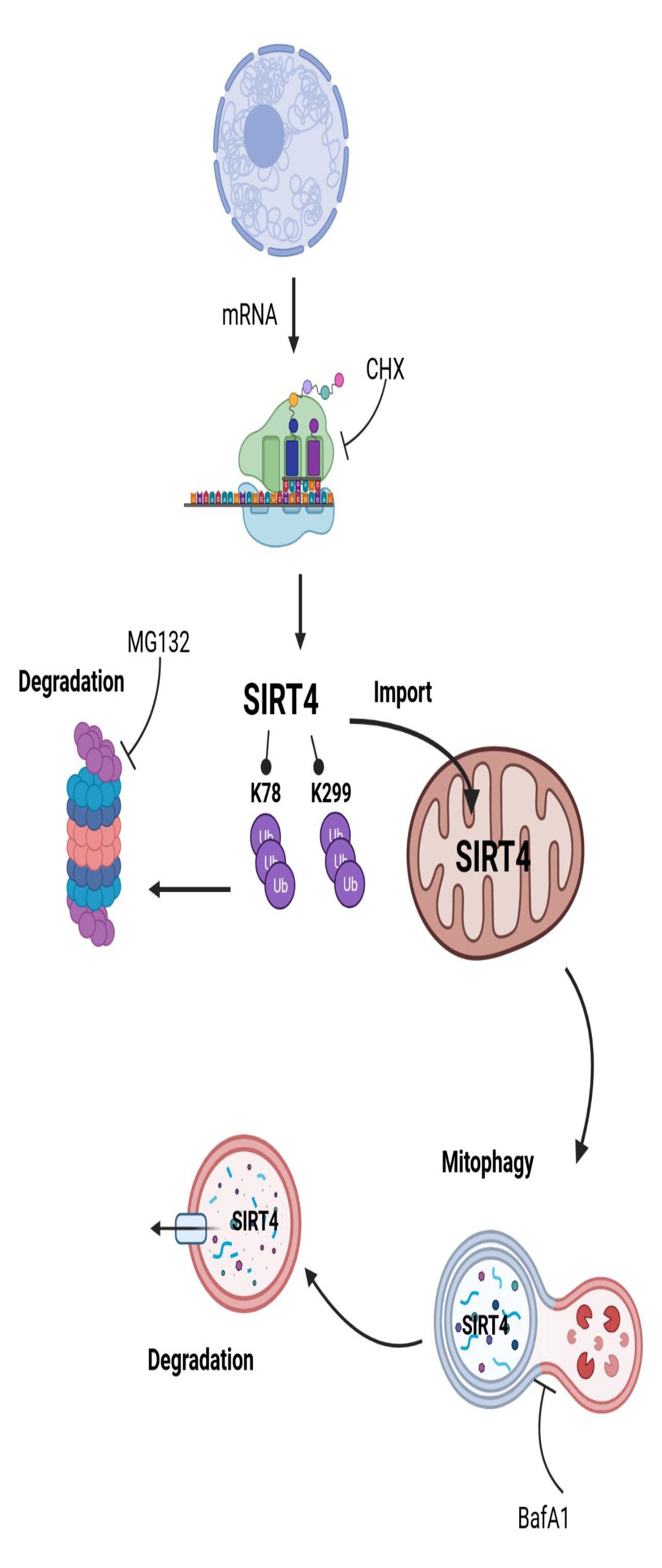
Model overview about the cellular pathways involved in protein degradation of SIRT4. Given the significantly increased protein half-life of SIRT4(K78R) (Figure 5C) and the greater stability of SIRT4(K299R) under pseudohypoxic stress (Figure 3), it is tempting to speculate that K78 and K299 play some divergent roles in basal *vs*. stress-induced degradation of SIRT4, respectively.

## Author Contributions

N.H. and R.P.P. initiated the project and designed the study. N.H., J.G., M.M., A.L., D.M.F., and R.P.P. designed, performed and analyzed the experiments. J.S., M.R.A, and D.M.F. provided expertise, tools, and essential reagents for mutational and nanobody-based co-immunoprecipitation analysis. N.H. and R.P.P. wrote the manuscript. All authors read, discussed, critically corrected, and approved the final version of the manuscript.

## Funding

This work was funded in part by the Stiftung für Altersforschung (grant 701.810.783) of the Heinrich Heine-University Düsseldorf (to R.P.P.), and by the Deutsche Forschungsgemeinschaft (DFG, German Research Foundation) project AH 92/8-3 (to M.R.A.).

## Acknowledgments

We thank Ursula Duerkop and Yvonne Arlt for expert technical assistance, Björn Stork for providing tools for polyubiquitination analysis, Sebastian Krüger for help with flow cytometry, Natascia Ventura for advice regarding the use of CoCl_2_ in pseudohypoxia, and Laura Bergmann for discussion.

## Conflicts of Interest

The authors declare no conflict of interest.

## Supplementary Figures

**Figure S1.**
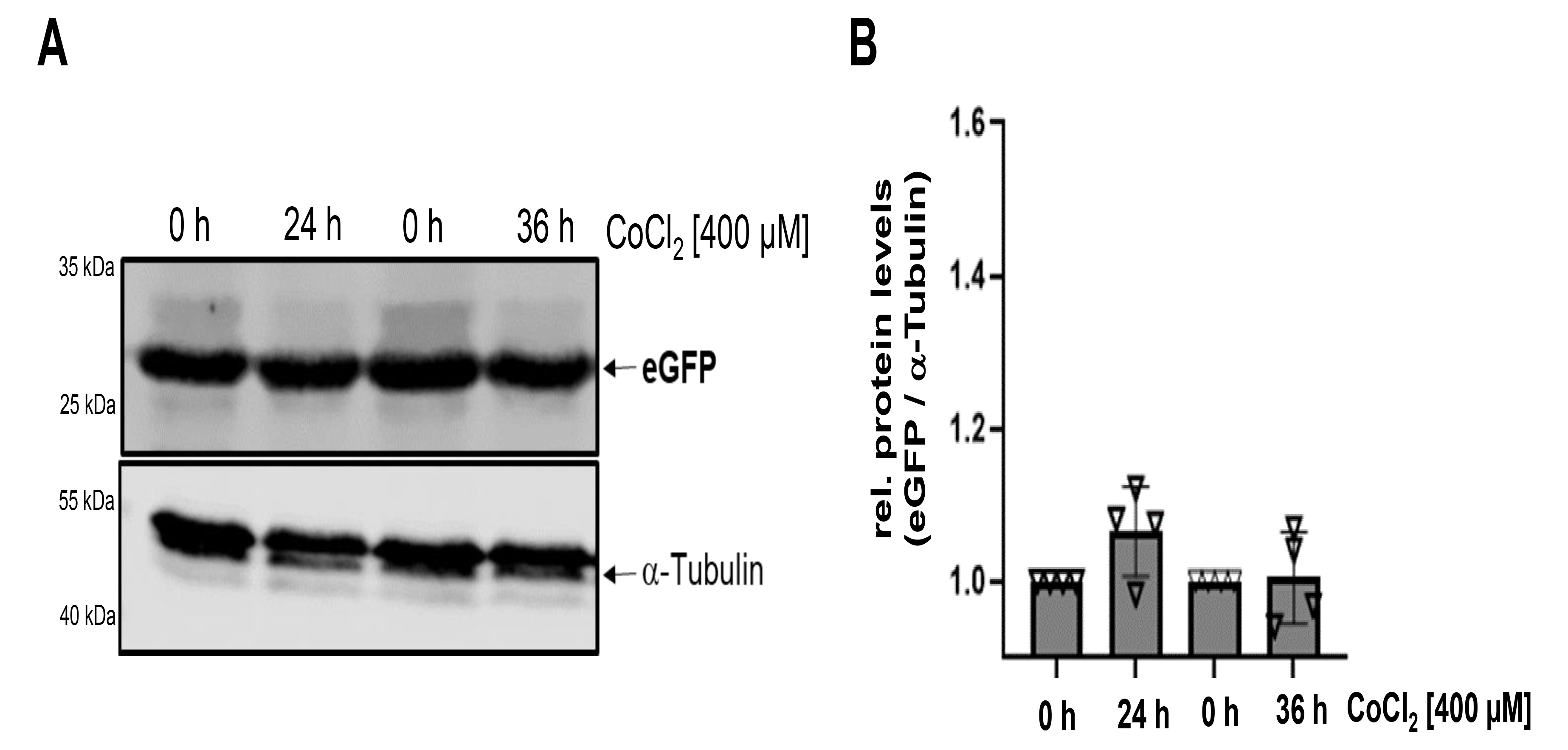
CoCl_2_ treatment of HEK293-eGFP cells does not lead to downregulation of eGFP levels. HEK293 cells stably expressing eGFP were subjected to CoCl_2_ treatment for 24 h and 36 h followed by immunoblot analysis. Relative quantification of eGFP immunoblot signals was performed using ImageJ based densitometric evaluation and α-Tubulin levels as loading control. To test statistical significances, a One-Way ANOVA test followed by Tukeýs Test was employed (mean ± S.D.).

**Figure S2.**
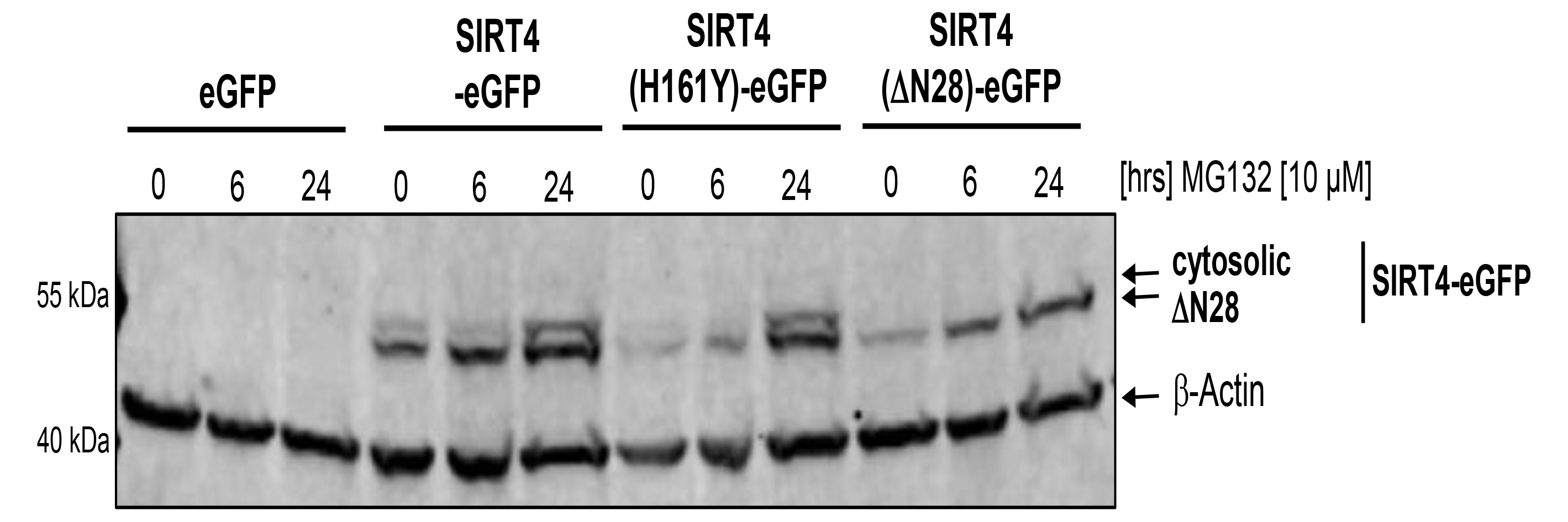
Stabilization of SIRT4(H161Y) and SIRT4(ΔN28) mutants by treatment with the proteasome inhibitor MG132. HEK293 cell lines expressing eGFP, SIRT4-eGFP, the catalytically inactive mutant H161Y, or the N-terminal deletion mutant SIRT4(ΔN28) that does not translocate into mitochondria, were subjected to MG132 treatment for 6 h and 24 h followed by immunoblot analysis of the respective SIRT4-eGFP/mutant SIRT4-eGFP levels. β-Actin staining served as loading control.

**Figure S3.**
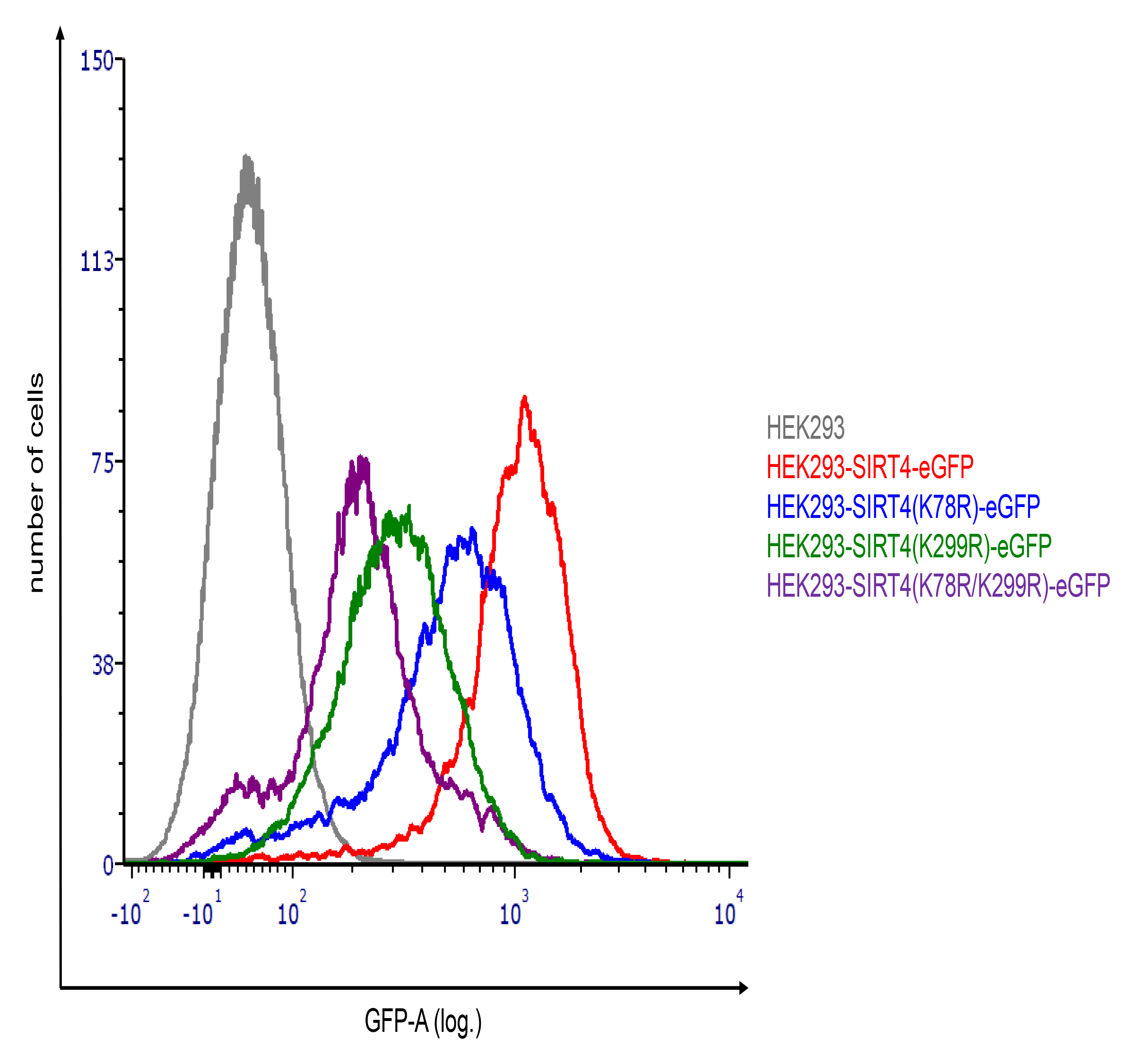
Flow cytometry-based analysis of expression of SIRT4 and SIRT4 mutants. Parental HEK293 cells as negative control and HEK293 cell lines stably expressing SIRT4-eGFP or SIRT4 mutants thereof were subjected to flow cytometry-based (GFP-A) expression analysis.

## Notes

### Competing Interest Statement

The authors have declared no competing interest.

